# Assessing the potential for assisted gene flow using past introduction of Norway spruce in Southern Sweden: Local adaptation and genetic basis of quantitative traits in trees

**DOI:** 10.1101/481614

**Authors:** Pascal Milesi, Mats Berlin, Jun Chen, Marion Orsucci, Lili Li, Gunnar Jansson, Bo Karlsson, Martin Lascoux

## Abstract

Norway spruce (*Picea abies*) is a dominant conifer species of major economic importance in Northern Europe. Extensive breeding programs were established to improve phenotypic traits of interest. In southern Sweden seeds used to create progeny tests were collected on about 3000 trees of outstanding phenotype (“plus” trees) across the region. Some were of local origin but many were recent introductions from the rest of the natural range. The mixed origin of the trees together with partial sequencing of the exome of >1,500 of these trees and phenotypic data retrieved from the Swedish breeding program offered us a unique opportunity to dissect the genetic basis of local adaptation of three quantitative traits (*height*, *diameter* and *budburst*). Through a combination of multivariate analyses and genome-wide association studies, we showed that there was a very strong effect of geographical origin on growth (height and diameter) and phenology (budburst) with trees from southern origins outperforming local provenances. Association studies also indicated that growth traits were highly polygenic and budburst somewhat less. Hence, our results suggest that assisted gene flow and genomic selection approaches could help alleviating the effect of climate change on *P. abies* breeding programs in Sweden.

## Introduction

Climate change is rapidly altering the environment of plants and animals, especially at high latitudes (Walther et al. 2002; Root et al. 2003). In order to alleviate the impact of climate change Aitken and Whitlock (2013) proposed to use assisted gene flow. The basic idea is that species currently adapted to dry and warm environments will be pre-adapted to the new environmental conditions prevailing in regions that experienced colder and wetter climates until today. Hence, facilitating introgression of alleles from Southern populations into more Northern ones could accelerate the process of local adaptation to the new climatic conditions.

Transferring material from Southern latitudes to more northern ones is not a new idea and extensive seeds transfer already took place in the past. Indeed, since the 1950s Sweden, Norway, and to a lesser extent Finland, started to import seeds of Norway spruce (*P. abies*), for forest reproduction material from Belarus, Czech Republic, Romania, Germany, Slovakia and the Baltic States (Myking et al. 2016; Jansen et al. 2017). As a matter of fact, we recently showed using genomic data that a very large part of the trees used to establish the Norway spruce breeding program in Southern Sweden corresponded to recent introductions (Chen et al. 2018). The aim of these introductions was twofold: (i) to obtain a large amount of seeds and (ii) to take advantage from the fact that trees from lower latitudes had a longer growth period and thereby a higher growth rate than local provenances when moved northwards (Dormling et al. 1968; Ekberg et al. 1979; Clapham et al. 1998).

More generally, transfer of trees within a range of 4 degrees of latitude was recommended to increase forest productivity (Persson and Persson 1992; Rosvall et al. 1998) but transfer from farther provenance could, in contrast, lead to maladaptation (Savolainen et al. 2013 and references therein). Indeed, the latter could be due to soil content, differences in biotic communities or to a mismatch between growth rhythm and optimal climatic conditions; for instance the risk of late-spring or early-fall frost damages increases with the range of the transfer.

In the present study we will take advantage of these extensive past transfers and their use in the breeding program to (i) assess the level of local adaptation of trees currently growing in Southern Sweden and (ii) investigate quantitative trait genetic architecture. The exome of more than 1500 trees that were used to create the Norway spruce breeding program for Southern Sweden were sequenced (Fig.1). Offspring of these trees were used in progeny tests across Southern Sweden in order to estimate the breeding values of their parents for three phenotypic traits of economic interest, *diameter*, *height* and *bud burst*. While *diameter* and *height* are related to wood production, *bud burst* reflects growth rhythm.

First, we showed a strong level of past local adaptation with high congruence between the clustering of genotypic, phenotypic and climatic data. Level of productivity of Southern provenances under current climatic conditions in Southern Sweden are thus higher because of a longer growth period. Through genome-wide association studies (GWAS), we then investigated both, the genetic basis of Norway spruce local adaptation to climate conditions and the genetic determinism of three phenotypic traits of economic interest. We identified a large number of genes involved in the response to environment variation or in the control of quantitative traits. Especially, we showed that the determinism of growth traits is much more polygenic than that of *bud burst*. Our data also highlight how traits with different pattern of geographical variation can be used to assess the impact of correction for population genetic structure in genome-wide association studies (GWAS). More importantly, we argued that while data from breeding programs might sometimes be incomplete or suboptimal, they are readily available and contains a lot of valuable information for evolutionary biologists.

## Material and Methods

### Trees sampling

The original sampling included 1672 samples from three related spruce species, Norway (*P. abies*), Siberian (*P. obovata*) and Serbian (*P. omorika*) spruces (Chen et al. 2018). In the present study, only 1545 trees from *P. abies* populations were considered (Fig. 1 and Tab. S1). These samples came from two sampling schemes:

i. 1475 individuals were “plus trees”, *i.e.* trees selected on the basis of their outstanding phenotype to create the base population of the Norway spruce Swedish breeding program. Needles were collected on trees from Skogforsk’s (The Forestry Research Institute of Sweden) clonal archives. Their progeny were represented in several progeny tests across Central and Southern Sweden (Fig. 1, black squares). Among 1475 “plus trees”, 560 had no clear records of their geographical origin (*i.e.* information on the origin of trees was missing in the archives of the breeding program, Tab. S1).
ii. 70 individuals were sampled from *P. abies* natural populations covering the main genetic domains of its distribution range (Lagercrantz and Ryman 1990; Tollefsrud et al. 2009; Tsuda et al. 2017; Chen et al. 2018). They were used as reference when defining the origin of the 560 “plus trees” whose origin was missing (Tab. S1).

**Figure 1:**
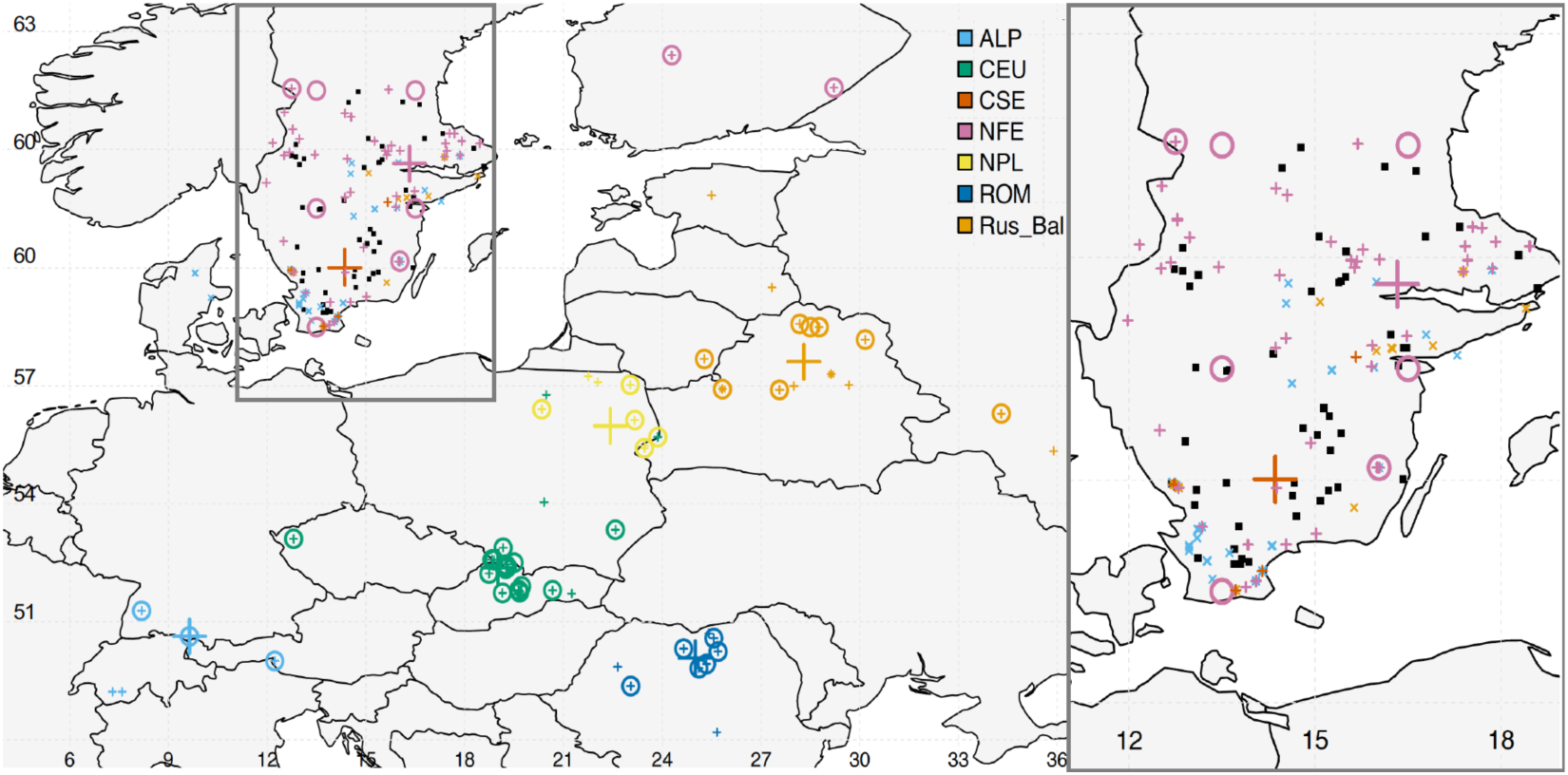
Trees original locations. Black squares are Norway spruce progeny test belonging to the Swedish breeding program. Small “plus” and “multiply” signs are tree sampling locations, the latter indicating a wrong assignation in available records. Large “plus” signs are the barycenter of the geographical coordinates of trees belonging to a same genetic cluster and having a known origin. Circles correspond to the centroid of the geographical coordinates of trees belonging to a same population as defined for SNP-environmental association analysis (see Methods § “SNP-environment relationships”). Colors correspond to genetic clusters (Carpathian, dark blue, ROM; Alpine, light blue, ALP; Central Europe, green, CEU; Northern Poland, yellow, NPL; Russia-Baltic, orange, Rus-Bal; Central and Southern Sweden, red, CSE; Fennoscandian, pink, NFE).

### Population structure analyses

The SNP dataset defined by Chen et al. (2018) was used to assess population structure and to define genetic clusters of the 1545 *P. abies* individuals.

#### SNP definition

The complete procedure for SNP identification is detailed in Chen et al. (2018). Briefly, after genomic DNA extraction, 40,018 probes of 20bp long were designed to cover exons of 26,219 *P. abies* contigs (Vidalis et al. 2018). Paired-end short reads were aligned to the reference genome of *P. abies* (Nystedt et al. 2013) using BWA-mem (Li and Durbin 2009). PCR duplicates were removed using PICARD v1.141 (http://broadinstitute.github.io/picard) and INDEL realignment was performed using GATK (McKenna et al. 2010) IndelRealigner. SNP calling was carried out using GATK HaplotypeCaller across all samples. After variant recalibration and filtering (≤ 2 reads for both, reference and alternative alleles and ≤ 50% coverage across individuals) 1,004,742 SNPs were retained. Finally, 917,107 *P. abies* SNPs were extracted for this study.

#### Genetic clusters definitions, inference of individuals’ unknown origin and relatedness

EIGENSOFT v6.1.4 (Galinsky et al. 2016) was used to perform principal component analysis (PCA) on the genetic variation of *P. abies* and to define subsequent genetic clusters based on 399,801 unlinked non-coding SNPs (pairwise LD ≤ 0.2 and FDR value ≥ 0.05, after haplotype phasing using MACH v 1.0, Li et al. 2010). Geographic origin of the 560 individuals for which no confident records of geographical origin were available was then assessed based on their genotype similarity to ascertained individuals. *P. abies* individuals of known origin were first grouped into seven major clusters based on genetic clustering results and their origin. These individuals were then used as a training set in a “Random Forest” regression model. The first five components of the PCA analysis were used for model fitting to classify the unknown individuals into each of the seven clusters. Five-fold cross-validation was performed for error estimation. Individuals of “unknown” origin were then assigned to the various genetic clusters defined from individuals from known origin. The whole regression process was repeated 1,000 times in order to estimate the confidence of each assignment.

Finally, for genome wide association studies, the individual relatedness (kinship) matrix was estimated using the “Centered IBS” method (Endelman and Jannink 2012) implemented in the TASSEL software (v.5.2.38, Bradbury et al. 2007).

### Phenotypes

Breeding values (BVs) for two growth traits, *diameter* and *height* and measures of *budburst* were extracted from the records of Skogforsk for 763, 808 and 834 “plus-trees”, respectively. Complete records of the three traits were available for 712 of those trees but the origin was known for only 283 of them. Briefly, for *height* and *diameter* more than 15 progenies of each “plus-tree” were planted in up to five progeny tests per tree scattered across Central and Southern Sweden. For *height* and *diameter*, the BVs were then computed, for each progeny test, using mixed linear model and BLUPs (Best Linear Unbiased Predictors) methodology through a restricted maximum likelihood approach from the following statistical model:

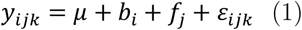

where, for a given trait, *y*_ijk’(ijk’(ijk’(_ is the observation for individual *y*_ijk’(ijk’(_ in block *y*_ijk’(ijk’(_, *y*_ijk’(ijk’(_ is the mean of the trait, *y*_ijk’(ijk’(_ is the fixed effect of block, *y*_ijk’(ijk’(_ is the random effect of family with a normal distribution 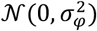 and *y*_ijk’_(_ijk’_) is the error term with a normal distribution 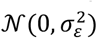. BVs were then reported as relative percentages to the mean. Therefore, 100 corresponds to the average BV and a relative BV of 110 thus indicates that the given genotype has a BV 10% higher than the average. For a given genotype, the average BV across the different progeny tests was then considered. *Budburst* was measured using Krutzsch scale (Krutzsch 1973), ranging from 0 (no burst) to 9 (full development of the needles) from a single clonal archive and transformed to normal scores based on midpoint values of the cumulative frequency distribution (Danell 1991) before analysis.

#### Inference of missing phenotype

Missing phenotypes (782, 737 and 711 respectively for *diameter*, *height* and *budburst*) were inferred from genotypic data using the “genomic selection” method implemented in TASSEL software (v.5.2.38 Bradbury et al. 2007; Zhang et al. 2010). Briefly, each trait was considered independently and the missing breeding values for a given trait were estimated using a mixed model that included a population structure matrix as fixed-effects and a kinship matrix as random effects to capture the covariance between genotypes. In other words, the BLUPs of individuals whose phenotype is missing are imputed from the phenotypes of closely-related individuals. Five-fold cross-validation was performed for accuracy estimation (20 iterations each).

### Relationships between phenotype and ancestral environment

The relationship between tree origins and their phenotypes was estimated using the following generalized linear model (GLM):

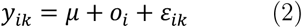

where, for a given trait, *y*_ijk’_(_ijk’_(is the observation for genotype *y*_ijk’_(_ijk’_((BVs for *height* and *diameter* or normal-score for *budburst*) from origin *y*_ijk’_(_ijk’_(, *y*_ijk’_(_ijk’_(is the mean value of the trait and *y*_ijk’_(_ijk’_(is the tree-origin specific fixed effect (a factor variable of seven levels corresponding to the seven genetic clusters) and *y*_ijk’_(_ijk’_(is the error term,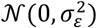 The significance of the difference between factor levels (genetic clusters) was computed from the complete model (2) using likelihood-ratio-test (LRT). Factor levels were grouped if no significant difference (*p* > 0.05) was identified and tree origin effect of reduced levels was further assessed as a simplified model (2).

#### Ancestral environment characterization

Data for 19 bioclimatic variables (monthly average for the 1970 – 2000 period, 10 arc minute resolution, ∼340 Km^2^) were downloaded from the online WorldClim database (v.2.0 http://worldclim.org/, Fick and Hijmans 2017). Two additional variables were computed from these data, the summer heat-moisture index (SHM) and annual heat-moisture index (AHM). A measure of yearly photoperiodic amplitude (*ΔDL*) was also computed as the difference between the average day length in June and the average day length in January (see table S2 for the complete list and details). For trees of unknown origin, bioclimatic data were extracted at the location corresponding to the centroid of the geographical coordinates of trees belonging to the same genetic cluster and having a known geographical origin (Fig. 1, large “plus” signs).

### Detecting loci underlying local adaptation

#### SNP-environment relationships

The Bayenv software (v.2, Coop et al. 2010; Günther and Coop 2013) was used to estimate correlations between allele frequencies at individual loci and bioclimatic variables, while accounting for population structure. A Bayesian mixed linear model, considering bioclimatic variables as fixed-effects and a variance-covariance matrix of allele frequencies as random-effect (to capture shared polymorphism due to populations common history) was fitted to population allele frequencies. In parallel, Spearman’s rank correlation coefficient, rho, was also computed from standardized allele frequencies, from which the covariance structure among populations was removed.

Forty-eight *P. abies* populations were defined by grouping trees from close geographic origins belonging to the same genetic cluster (≥ 5 trees per population, Fig. 1, circles). Note that only trees with a known origin were considered as the populations were defined from geographic coordinates (777 trees). Twenty variance-covariance matrices were estimated from 20,000 non-coding and unlinked SNPs randomly sampled from the 399,801 non-coding and unlinked SNP dataset. The average matrix across the 20 runs was then considered in the model. Finally, for each population, bioclimatic data for each tree location were averaged. For each climatic variable, the following filtering (based on Bayes Factor and Spearman’s rho) was applied to retain only the most relevant SNPs: *i)* the SNPs were ranked according to their Bayes Factor (BF) and a SNP was retained if its BF > 150 (very strong strength of evidence according to Kass and Raftery (1995) or, if its BF > 20 (strong strength of evidence) and was within the 0.1% highest BF; *ii)* in parallel SNPs were ranked according to Spearman’s rho and only those that were satisfying the first criteria and were within the 1% highest absolute rho were conserved for further analysis, as recommended by Günther and Coop (2013).

#### SNP-phenotype relationships

For each trait independently, the additive allelic effect of each SNP on the phenotype and the corresponding standard-error were estimated through a linear mixed model considering population structure (first three principal components of a SNP-based PCA) and individual relatedness (kinship matrix). The analysis was performed through compressed mixed linear model (Zhang et al. 2010) implemented in the R package GAPIT (Lipka et al. 2012). For this analysis, only bi-allelic SNPs with a minimum allele frequency higher than *y*_ijk’_(_ijk’_(and a minimum number of individuals per genotype of 10 were considered.

The statistical significance of the SNP associated to the three phenotypic traits was investigated using a recently developed Empirical Bayes approach for adaptive shrinkage (Stephens 2017) implemented in the R package *ashr* (Stephens et al. 2018). Traditional False Discovery Rate (FDR, Storey 2003) methods are based on the sole *p*-values. In contrast, *ashr* uses both allelic effect sizes and standard errors. It models the GWAS results as a mixture of SNPs that have a true effect size of exactly zero and SNPs that have a true effect size that differs from zero. The “local false sign rate”, *lfsr*, which refers to the probability of getting the sign of an effect wrong, is then used as a measure of significance and to compute *s*-values (Stephens 2017), which are the analogues of Storey’s *q*-values (Storey 2011). The “local false sign rate” is therefore more robust to errors in model fit than FDR (Stephens 2017).

#### Gene function and enrichment test

Gene ontology (GO) enrichment was performed using the “top GO” R package (v2.26.0; Alexa and Rahnenfuhrer 2010). Annotation from ConGenIE (the Conifer Genome Integrative Explorer, http://congenie.org/) was used as reference (*i.e.* custom input), all the GO terms were conserved (nodeSize parameter = 1). For the various lists of candidate genes defined through both SNP-environment and SNP-phenotype analyses, enrichment of genes in particular GO terms biological processes (BP) was assessed using the ‘weight’ algorithm and Fisher exact test (*p* < 0.05). Finally, the REViGO software (Supek et al. 2011) was used to remove GO terms redundancy and to cluster the remaining terms in a two dimensional space derived by applying multidimensional scaling to a matrix of the GO terms semantic similarities (default parameter setting: allowed similarity = 0.7, SimRel to measure the semantic similarity, UniProt as database). The Cytoscape software v3.6.1 (Shannon et al. 2003) was then used to visualize GO terms networks.

## Results

### The Norway spruce breeding program is genetically diverse and representative of the whole range

The Norway spruce breeding program in Southern Sweden was started in the fifties from plus-trees collected over a thirty-year period in Southern and Central Sweden. Because of the continuous introduction of material from the natural range of Norway spruce, the Norway spruce breeding program today includes individuals from the five major geographic domains: the Russia-Baltics region (Russia, Belarus, Estonia, Latvia, and Lithuania), the Visegrad group (Hungary, Poland, Czech Republic, and Slovakia), the Fennoscandia (Norway, Sweden and Finland), the Alpine domain (France, Italy, Switzerland, Germany and Austria) and the Carpathian domain (mainly Romania). The extent to which each of these groups contributed to the breeding population was however unclear because the information on the origin of around one third of the individuals was missing. In a previous study (Chen et al. 2018), based on ∼400K non-coding and unlinked SNPs, we were able to assign each tree to one of the seven *P. abies* genetic clusters: four of these clusters corresponded to the major domain described above Alpine (hereafter, ALP), Carpathian (CAR), Fennoscandian (FEN) and Russia-Baltics (Rus-Bal). However, the Visegrad region actually comprised two genetically distinct clusters, Central Europe (CEU, Hungary, Czech Republic, Southern-Poland, and Slovakia) and Northern-Poland, due to introgression of *P. obovata* genetic background into the latter. The last one, in Central and Southern Sweden (CSE) appeared from the hybridization of trees belonging to ALP and FEN groups (for more details, please refer to Chen et al., 2018). These analyses thus allowed us to identify the origin of 560 trees used in the breeding program but also to correct the origin of 134 trees that had ambiguous record (“x”symbols in Fig 1 and Fig. 2A).

**Figure 2:**
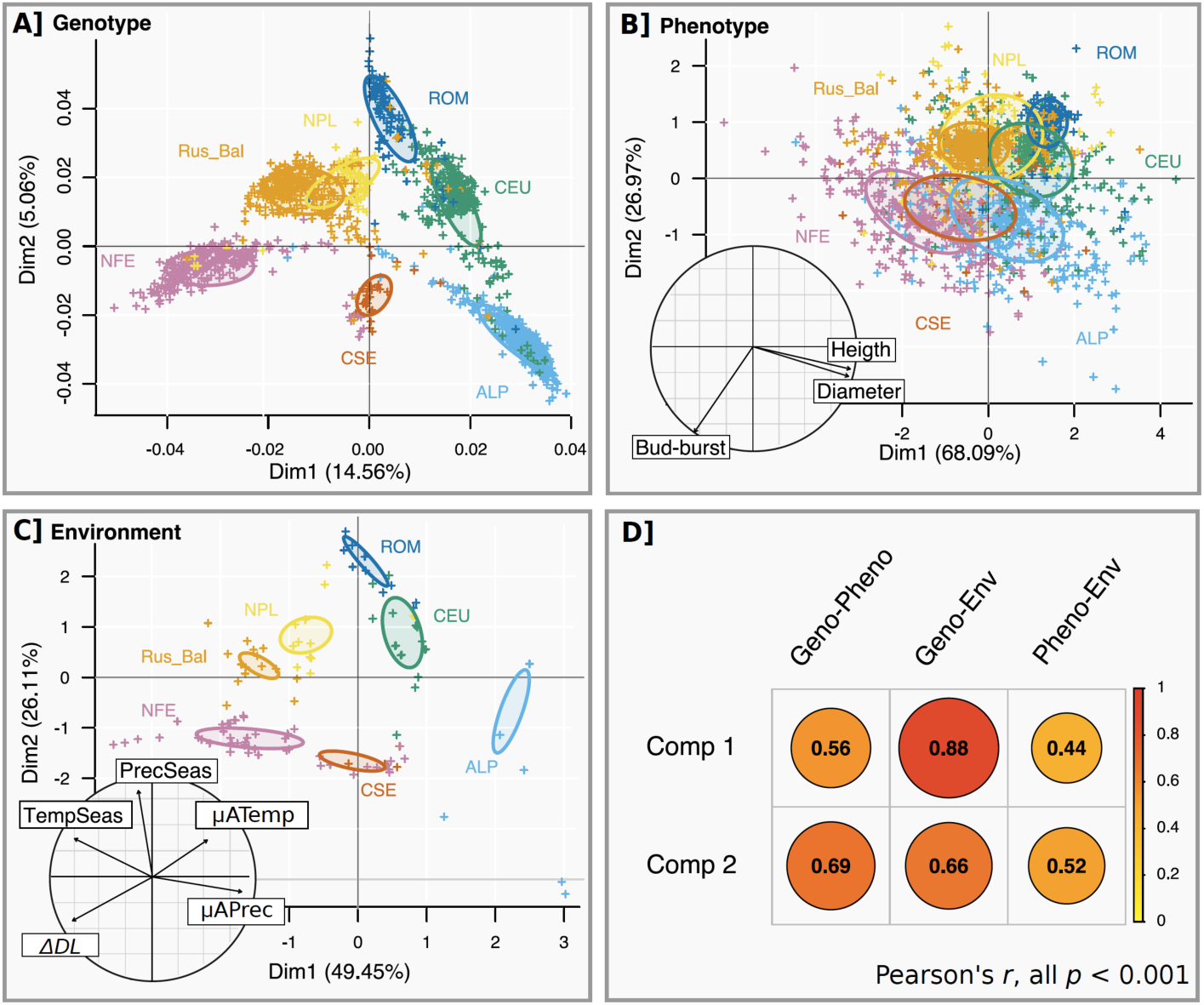
Pattern of variation of trees genotype, phenotype and original environment. Principal component analyses (PCA) based on **A**, SNPs data (modified from Chen et al. 2018), **B** Phenotypic data and **C**, climatic variables of the populations of origin (see Tab. S2 for climatic variable details). Colors correspond to genetic clusters (Carpathian, dark blue, ROM; Alpine, light blue, ALP; Central Europe, green, CEU; Northern Poland, yellow, NPL; Russia-Baltic, orange, Rus-Bal; Central and Southern Sweden, red, CSE; Fennoscandian, pink, NFE). Panel **D** represents Pearson’s product moment correlation coefficient, *r*, between principal component 1 (Comp 1) or 2 (Comp 2) of the various PCA, disc diameters are proportional to the corresponding correlation coefficient.

### Ancestral environment is a strong predictor of phenotype in Norway spruce

Trait values estimated from progeny tests across Southern Sweden were then used to assess the influence of tree origin (genetic cluster) on phenotypes. All three traits, *diameter*, *height* and *budburst*, differ among genetic clusters (Model 2, *F* = 39, *df* = 6, *p* < 0.001; *F* = 20, *df* = 6, *p* < 0.001; *F* = 12, *df* = 6, *p* < 0.001; respectively for *diameter*, *height* and *bud burst* Fig. S1).

Climatic variables at tree original locations were then used to characterize the environment at origin (see table S2 for more details) and investigate environment-phenotype relationships. Unfortunately, due to missing records, tree origin and phenotype information were both available for only 279 trees. Both datasets were thus completed by considering bioclimatic data at the location corresponding to the centroid of the geographical coordinates of trees belonging to the same genetic cluster and having a known origin, (Fig. 1, large “plus” sign), and by imputing missing phenotypes using linear mixed models (see Methods, § “Missing phenotype inference”). Correlations between observed and predicted phenotypes were strong (Spearman’s *y*_ijk’_(_ijk’_(= 0.99 for *diameter* and *height* and *y*_ijk’_(_ijk’_(= 0.83 for *budburst*, Fig. S2) but accuracy was much higher for *diameter* and *height* (0.52 and 0.41, respectively) than for *budburst* (0.27). This provided us with a complete dataset for the 1543 trees.

Genotypic (∼400K SNPs), phenotypic (*diameter*, *height* and *budburst*) and environmental variables were each then described through principal component analysis (PCA, “ade4” R package *v.1.7-10*, Chessel et al. 2004). The latter were characterized by annual mean and seasonality of precipitation (μAPrec and PrecSeas, resp.) and of temperature (μATemp and PrecTemp, resp.), as well as by an indicator of photoperiod (*Δ*DL; see Tab. S2 for more details). Strikingly, genotype, phenotype and environment data presented very similar clustering patterns (Fig. 2 A, B and C) and the principal component coefficients of the different PCA were highly correlated (Fig. 2D).

Further, a more thorough investigation of the phenotype-environment relationships (21 climatic variables considered, see complete list in Tab. S2) showed that *diameter* and *height* decreased mainly along a South to North latitudinal gradient (Pearson’s *r* = −0.62 and −0.47, respectively, all *p* < 0.001) and were thus strongly associated to climatic variables following this gradient (e.g., *ΔDL*: *r* = −0.61 and −0.46; summer heat-moisture index, SHM: *r* = - 0.54 and −0.38; annual precipitation, μAPrec: *r* = 0.50 and 0.31; all *p* < 0.001 Tab. S2 and Fig. S3, for the complete analysis). On the other hand, *budburst* followed both a latitudinal gradient (South to North, *r* = 0.39, *p* < 0.001) and a longitudinal gradient (West to East, *r* = −0.47, *p* < 0.001). It was thus more associated with climatic variables reflecting continentality (e.g., PrecSeas: *r* = −0.54; mean diurnal temperature range, μRangeDuir: *r* = −0.52; annual temperature range, ARangeTemp: *r* = −0.39; all *p* < 0.001. See Tab. S2 and Fig. S3 for the complete analysis).

### Association mapping revealed close link to environment of origin despite strong population structure

In order to identify loci underlying local adaptation in Norway spruce, different populations were defined based on genetic clusters and tree original locations (*i.e*. trees from close geographic origins belonging to the same genetic cluster were grouped, Fig. 1, “circles”); note that only trees with a known origin were used in the present analysis (48 populations, 777 trees, Tabs. S3). Climatic variables were then used as fixed factors in independent mixed linear models (MLMs) accounting for population structure (Bayenv software v.2, Coop et al. 2010; Günther and Coop 2013), to explain SNP frequencies variation across populations (Fig. S4).

After stringent filtering steps to control for false positives (see Method § “SNP-Environment relationships”), many SNPs were found to be associated with environmental variables (min = 46 for Temp_1, annual mean temperature and max = 344 for Prec_4, total precipitation of the driest month, Tab. S4). Approximately 25% of these SNPs belong to intergenic regions, a half belongs to introns and the remaining quarter belongs to exons (Tab. S4; SNP annotations and transcript descriptions are given in supplementary material 1). For each climatic variable, the redundancy varied a lot from 0 to ∼65%, meaning, for the latter, that two third of the candidate SNPs are related to the same transcripts (Tab. S4). Despite such a degree of redundancy, many genes were found to be associated with each climatic variable (min = 43 for Temp_1, annual mean temperature, and max = 148 for photoperiod, Tab. S4). On average, a larger number of transcripts were associated with precipitation related variables (*y*_ijk’_(_ijk’_(, mean overlap between variables 44%, Tab. S5A) than with temperature related variables (*y*_ijk’_(_ijk’_(, overlap 12%, Welch’s two-sample *t*-test, *t* = −4.9, *df* = 11, *p* < 0.001) or moisture (*y*_ijk’_(_ijk’_(, overlap 38%). The largest group consisting of transcripts associated to photoperiod (*y*_ijk’_(_ijk’_(Tab. S4). Importantly, the present study demonstrates that climatic variation creates a widespread selective pressure in the Norway spruce genome as non-overlapping categories of genes were associated to different climatic variables. For instance, the average overlap between genes involved in response to temperature variables and those involved in response to precipitation was only 5% (Tab. S5B).

However, this limited overlap at the gene level, masks a much more important overlap at the functional level as many of these genes tend to be involved in the same biological processes as shown by the strong overlap between gene ontology terms (supplementary material 1). The smallest overlap was between photoperiod and moisture index, 15%, and the largest, 72%, expectedly, was for precipitation and moisture (Fig. 3A and Fig. S5A).

**Figure 3:**
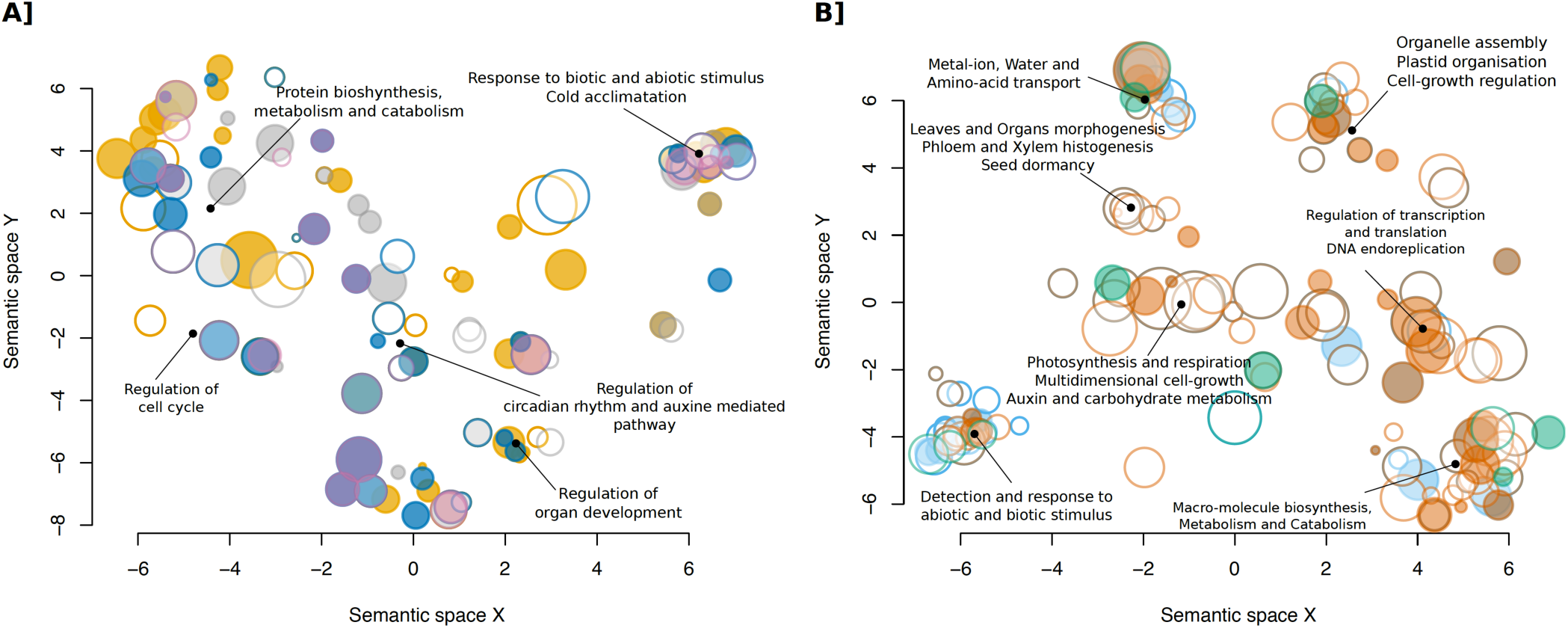
Two-dimensional semantic space representation of biological processes gene ontology terms. Semantic representation of GO categories are colored according to a significant over-representation (Fisher’s exact test < 0.05) within genes showing significant associations with climatic variables (**A**, *temperature* in orange, *precipitation* in blue, *moisture* in pink and *photoperiod* in grey) or involved in the determinism of phenotypic traits (**B**, *diameter* in blue, *height* in brown and *bud-burst* in green). The circle diameter is proportional to the number of aggregated GO terms. Descriptions of the main GO term categories are provided (see Fig. S4 and S5 for the complete annotation).

**Table 1:** Linear regressions between either trait values or between allelic effect sizes.

| Trait A | Trait B | Phenotypic |  |  | All SNPs |  |  | A significant SNPs |  |  | B significant SNPs |  |  |
| --- | --- | --- | --- | --- | --- | --- | --- | --- | --- | --- | --- | --- | --- |
| | | $r^2$ | $t$ | $df$ | $r^2$ | $t$ | $df$ | $r^2$ | $t$ | $df$ | $r^2$ | $t$ | $df$ |
| Height | Bud-burst | 0.12 | -15 | 1553 | 0.01 | -43 | 161112 | 0.12 | -5 | 173 | 0.9 | -16 | 28 |
| Diameter | Bud-burst | 0.08 | -12 | 1553 | 0 | -26 | 161112 | 0.01 | -2<sup>ns</sup> | 178 | 0.9 | -16 | 28 |
| Height | Diameter | 0.72 | 63 | 1553 | 0.64 | 539 | 161112 | 0.95 | 58 | 173 | 0.95 | 59 | 178 |

$r^2$ : Adjusted-R-squared; $t$ : t-statistic value, all are significantly different from 0, $p < 0.001$ unless indicated (ns); $df$ : degree of freedom.

### Growth traits and budburst have different genetic architecture

To characterize the genetic architecture of the different phenotypic traits and to identify SNPs involved in their determinism, a genome wide association studies (GWAS) was conducted with MLMs. Population structure was considered by including the three first principal components of a SNP-based PCA as fixed effects and a kinship matrix as random effect in the MLM; note also that only trees with a measured phenotype (*i.e.* not inferred) were included in these analyses (763, 808 and 834 trees, respectively for *diameter*, *height* and *budburst*).

*Diameter* and *height* had a highly polygenic determinism as no less than 180 and 175 SNPs, respectively, had a significant effect on trait values (*s*-value < 0.1, *s* is the analogue of *q*-value for *false sign rate* detection, see Methods §“SNP-Phenotype relationships”, Tab. S4 and Fig. 4A). In striking contrast, a mere 32 SNPs were detected for *budburst*. These SNPs affected more than 130 different genes for growth traits (∼20% redundancy) but only 15 genes for *budburst* (∼50% redundancy).

**Figure 4:**
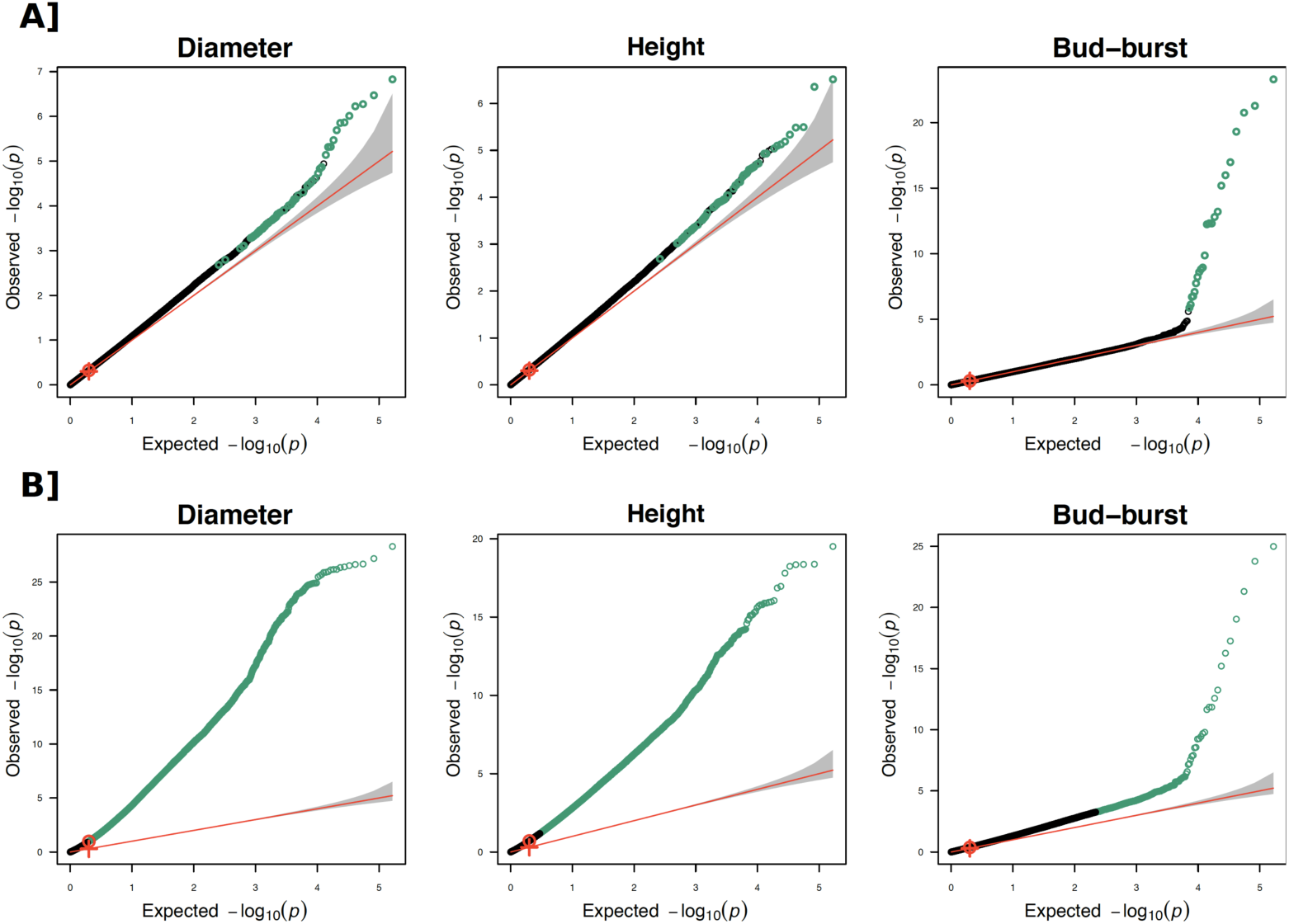
Q-Q plot of observed *vs* expected *p*-values. For each phenotypic trait, −log_10_ *p*-values, controlling for population structure (**A**) or not (**B**), are represented as a function of expected −log_10_ *p*-values assuming a uniform distribution. The red line is the one to one quantile line and the grey area is the 95% confidence intervals around it. The red circle and the “plus” sign represent the medians of observed and expected *p*-values, respectively. Significant corrected *p*-values (*s*-value < 0.1, see text) are colored in green.

Moreover, the phenotypic correlations between traits are due to pleiotropic effects at the genotypic level (Tab. 1 and Fig. S6). For instance, tree height and diameter were strongly correlated (Spearman’s *rho* = 0.86) and such a correlation was due to correlation between SNP allelic effect sizes (*rho* = *0.91* and *0.93*, when considering SNPs significant for *height* or SNPs significant for *diameter*, respectively). The pattern for *budburst* was different as SNPs involved in its control had a strong influence on both *height* and *diameter* (*rho = −0.92* and *rho = −0.91*, respectively), but the converse was not true (*rho = - 0.36* and *rho = −0.05*).

Finally, as for SNP related to environment, genes associated to the variation in phenotypic traits belong to biological processes, functions, pathways and network expected for the trait under consideration. For instance, genes involved in the control of phenotypic traits are associated to GO terms such as, to name a few, regulation of auxin metabolism, response to light and photoperiodism, gravitropism, cell growth or organs development (supplementary file 1). Furthermore, in contrast with climatic variables, at the network scale, GO terms associated only with *diameter* and those associated only with *height* tended to cluster (Fig. 3B and Fig. S5B).

## Discussion

Norway spruce (*Picea abies*) is a dominant conifer species of major economic importance in Northern Europe. Extensive breeding programs were established to improve phenotypic traits of interest, focusing on productivity, wood quality and resistance to pathogens. Here, genetic and phenotypic information was collected on more than 1500 trees of outstanding phenotype (“plus” trees) that were used to establish the breeding population. Some of these trees were of local origin but many corresponded to recent introductions from the rest of the natural range. In the present study we demonstrated that these data present a unique opportunity to study the genetic basis and the role of local adaptation in the determinism of quantitative traits. This last point is crucial for breeders and forest managers with a mean to assess the potential of assisted migration as a strategy to mitigate the impact of climate change on forest productivity and health.

### Caveats and solutions

We used breeding values for two phenotypic traits, *height* and *diameter* that were collected from different series of progeny tests of the Swedish breeding program. While it is unquestionable that breeding programs are a treasure trove for biologists, working from data originating from series of trials planted in the early 80s includes some serious challenges.

First, the data are heterogeneous. Indeed, the phenotypic data were collected on trees that were planted in progeny tests located at different latitudes in Sweden and the age of the trees at measurement varied across trials (from 6 to 15 years old for *height* and from 9 to 15 for *diameter*). A part of that variance was considered when computing the breeding values within trials as trees from the same trial were of the same age and obviously faced the same environment, but this nonetheless neglected genotype by environment interactions and did not remove the age variation across trials. In some trials breeding values for height were computed with five years intervals and were highly similar (*r^2^* ∼ 0.8, data not shown) and genotype by environment (GxE) interactions are known to be weak in *P. abies* breeding program in Central and Southern Sweden (Berlin et al. 2015). The heterogeneity introduced by these two effects should thus be somehow limited.

Finally, the trees belonged to different trial series, a trial series being a set of progeny tests comprising the same individuals. If the average BVs across a trial series is used, then GxE interactions are to some extent included. However, it should be pointed out that BVs from different trial series are not strictly speaking comparable as they were analyzed separately. As the BVs from each trial series were compiled and used as one complete data set there may be a bias. Such a bias would have been avoided if all the trial series had been evaluated simultaneously with e.g. the TREEPLAN system (McRae et al. 2003). The BVs would then have been truly comparable on the same scale but this was unfortunately not possible at the time the present study was initiated. Interestingly, while this heterogeneity will certainly have weakened the clustering of the individuals based on their phenotypic data it did not erase it altogether. Hence, our results are conservative.

Second, the data are incomplete: for roughly a third of the dataset the exact coordinates of the origin of some of the samples were unknown and phenotypic data were missing for about half of the trees. These difficulties were circumvented by using a large scale SNP dataset and supervised machine-learning algorithm to precisely assign each genotype to a given geographic origin (accuracy > 0.92, see Chen et al. 2018) and to infer the phenotypic values for trees for which records were lacking (genomic selection). Despite heterogeneity in the phenotypic data the method was accurate enough (e.g. > 50% for *diameter*) to provide us with a complete dataset (∼1500 trees) for studying phenotype – environment relationships.

Re-sequencing technologies are continuously developed and their efficiency keeps increasing. They are now affordable (and prices are still decreasing) and it is now possible to obtain genomic data for a large number of individuals. Thanks to new statistical approaches, mostly based on machine learning (see Schrider and Kern 2018, and examples within), it is now possible to overcome issues often encountered with such large and long-term survey datasets such as incompleteness. Breeding programs thus represent a valuable and still underused source of study material for evolutionary biologists. This is especially true for forest trees, as progeny tests and common gardens require extensive space and need to be measured over long periods of time, something that cannot easily be done today within universities or research institutes; in our case, the trials were spread all over Southern Sweden and some were started half a century ago.

### Population structure, local adaptation and genetic architecture of quantitative traits

Animal and plant species are known to have undergone cycles of contractions and expansions as a result of successive glacial and interglacial periods (Bennett 1997). The contraction phase is responsible for reproductive isolation between refugial area, which, in association to bottlenecks (reduction of the effective size of a population), can lead to a strong divergence between populations or even speciation events (Petit et al. 2003). During the expansion phase, a secondary contacts between the genetic entities can occur, resulting in introgression that can play a major role in the evolution of species (see Arnold 2004 and references within). However, (re)-colonization also involves facing environmental changes and natural selection also played a role in the current distribution of species (Saccheri and Hanski 2006). The phenotypes of individuals are therefore the result of the complex interplay of the demographic history of populations and of local adaptation. However, because (re)-colonization routes often followed environmental gradients, these two effects are generally confounded and disentangling the role of each in trait evolution remains challenging (Gaggiotti et al. 2009).

In *P. abies*, contraction phases resulted in three strongly differentiated genetic clusters, a Northern domain in Fennoscandia and two Southern domains in the Alps and the Carpathians that have been amply documented (Borghetti et al. 1988; Lagercrantz and Ryman, 1990; Acheré et al. 2005; Heuertz et al. 2006; Tollefsrud et al. 2009; Tsuda et al. 2016; Chen et al. 2018). These three genetic clusters had a major impact on phenotypic divergence as illustrated in our study and in the seminal work of Lagercrantz and Ryman (1990). In the latter, the authors analyzed 48 Norway spruce provenances at both 22 allozyme loci and seven morphological characters describing seed, growth and phenology. They analyzed both allozyme and phenotypic variation with Principal Components Analysis and, as in our case, observed a striking similarity between the two resulting plots indicating that population history had a strong impact on the divergence of phenotypic traits. In our case, the fact that environmental variables at the locations of origin of the different clusters show the same clustering as both phenotypic traits and genetic polymorphism strongly suggests that this divergence is not entirely neutral and may reflect local adaptation (Savolainen et al. 2013). This agrees with other studies of local adaptation in forest trees that have all concluded that local adaptation is common in forest trees despite extensive gene flow (e.g. Chen et al. 2012, 2014; Avia et al. 2014; Lind et al. 2014; Yeaman et al. 2016). This apparent paradox was first explained by Le Corre & Kremer (2003) (see also Kremer and Le Corre 2012; Le Corre and Kremer 2012). Their model was later extended by Berg and Coop (2014) which, in brief, shows that high differentiation between populations at quantitative traits will not result from large change in allele frequencies at a limited number of loci but instead will follow from coordinated small changes in allele frequencies at a myriad of loci, each of small effect underlying the variation in the quantitative traits.

We indeed found that the three traits used in the present study were highly polygenic, albeit *height* and *diameter* appeared more polygenic than budburst. Incidentally, our results also have important consequences for the estimation of trait polygenicity and more specifically for understanding the presence of a large number of false positives due to population structure. Chen et al. (2018), indicated the presence of secondary contacts between these main domains (Fig. 2A) and current *P. abies* populations are mainly structured along a latitudinal gradient as are the climatic variables influencing growth traits (Fig. 2B and C). In contrast, *budburst* varies along both latitudinal and longitudinal gradients (Fig. 2B). A lower confounding effect of population structure is thus expected for *budburst* than growth traits when investigating trait genetic architecture. In order to evaluate the impact of population structure on our ability to detect growth-trait related SNPs we reproduced the GWAS but without controlling for population structure (Fig. 4). As expected for *budburst*, *p*-values were biased toward lower values, going down from 761 significant SNPs to 32, after correction for population structure and multiple testing. While significant and already rather massive, this effect was minor compared to the impact of population structure on SNPs associated to growth traits where the number of significant SNPs went from > 50,000 without correction for population structure to ∼180 with correction (Fig. 4). Obviously, population structure impeded a proper detection of SNPs affecting growth traits in *P. abies*, because of likely over-correction for population structure. It also means that our estimates of ∼180 SNPs affecting growth traits is likely to be conservative and thus that the genetic architecture of *height* and *diameter* is highly polygenic. These results are in line with what was recently described for other quantitative traits determinism in model species (e.g. Daub et al. 2013; Berg et al. 2017; Boyle et al. 2017), reconciliating genomic data with Fisher’s infinitesimal model (Barton et al. 2017; Turelli 2017).

### Southern genotypes outcompete local trees for growth traits

The strong association between genotype and phenotype variation showed that *height*, *diameter* and *budburst* possess a strong genetic determinism. On the other hand, the association between genetic diversity and the environment of origin reflected a strong influence of population evolutionary history on genetic diversity. Finally, the fact that phenotypic traits followed environmental gradients revealed a strong pattern of local adaptation of Norway spruce populations to their original environment. Yet, despite this strong signature of local adaptation to the home environment, Southern genotypes, for instance those from Romania, were taller and larger than Northern ones when grown in southern Sweden although they resumed growth later in the spring than most northerly provenances. By linking phenotypic data to climatic variables, our study, as several before it (Heide 1974; Dormling 1979), highlights the importance of temperature in the determinism of *budburst* and of, temperature, precipitation and photoperiod, on growth traits. Given climate change expectations in Southern and central Sweden (+2 to 4°C annual mean temperature and higher annual precipitations (Swedish commission on climate and vulnerability, 2007), it will be crucial to consider the tree origins in future development of the breeding program in these regions.

A major limitation for assisted gene flow for boreal species come from the risk of frost damages due to late-spring frost for Northern genotypes (because of a too early *bud break*) or early-fall frost for Southern ones (because of a too long growth period, e.g. Montwé et al. 2018 but see MacLachlan et al. 2017, 2018 for lodgepole and interior-spruce, respectively). In the present case, frost damages were recorded within nine trials (135 trees) and no difference in frost-resistance was observed between genetic clusters despite differences between trials (data not shown). In the long-term, episodic-frost are expected to decrease in Central and Southern Sweden given global warming (Swedish commission on climate and vulnerability, 2007); even if frost-damage risk could first increase during a transient period (Langvall 2011). Given the superior productivity of more southerly provenances, assisted migration appears as a strategy worth of further testing. In particular, biotic communities (insects, fungus, microbiome) and soil content represent two additional sources of maladaptation for large scale transfers that deserve further scrutiny (e.g., Wang and Klinka 1997; Macel et al. 2007; Aitken et al. 2008; Crémieux et al. 2008; Vitt et al. 2010). Indeed, in our different GWAS, we detected numerous genes involved in immune system responses as well as in metal-ion transport or pH-regulation. Investigating the relative impact of the afore-mentioned risks would require additional studies to get a complete assessment of the effect of local soil and biotic communities on non-local genotypes.

Finally, thanks to large scale genomic data (>400K SNPs), we were able to characterize *P. abies* population structure at a finer scale than in previous studies (Chen et al. 2018). This indicated the existence of a new genetic cluster corresponding to hybrids between Fennoscandian and Alpine trees (CSE cluster). In the present study, we further showed that trees belonging to that cluster also had an intermediate phenotype evidencing that if some limitations to Southern genotypes settlement exist they are clearly not strong enough to impede Alpine trees to reproduce with local ones. Hence, assisted gene flow would in the long-term lead to a dilution of the local gene pool, a risk that should certainly be considered.

## Conclusion

The sequencing of >1500 Norway spruce trees coming from the Swedish breeding program allowed us to analyze the influence of tree origins on phenotypic traits and to investigate their genetic basis. From a practical point of view, our study lends support on strategies based on assisted gene flow to alleviate the impact of climate change in Central and Southern Sweden breeding program. First, trees with Southern origin are taller and bigger than local ones (two valuable characteristics for wood industry), and second, we showed that the determinism of these traits is highly polygenic, arguing for genomic selection approaches for trait improvement, especially considering the strong genetic correlation between both traits. From a more general perspective, our study revealed a strong pattern of local adaptation in Norway spruce, phenotypic traits following environmental gradient and trees origin explaining a large part of their variance. It also showed that breeding programs are valuable resources for large scale genomic study. By strongly controlling environmental variance, they are ideal systems for trait-determinism investigation. Re-sequencing technologies being continuously developed and becoming more affordable, it will soon be possible to sequence a number of individuals large enough to apply statistical methods currently limited to humans and a handful of model species, allowing investigating, for instance, the strength and direction of selection acting on a trait of interest (Guo et al. 2018; Zeng et al. 2018 and references therein). Finally, our study showed that comparing traits that followed different geographic gradients could help to better comprehend and address the confounding effect of population structure on GWAS. Developing a statistical framework to control for population structure using this insight will, however, require further investigation.

## Supporting information

## Acknowledgements

The present study was financed by the Swedish Research Council for Environment, Agricultural Sciences and Spatial Planning (FORMAS) to ML, GJ and BK and by the Swedish Foundation for Strategic Research (SSF) project “Genomic selection of Norway spruce for new bioproducts”.

## Supporting Information

- **Table S1:** Trees origin from available records.
- **Figure S1:** Influence of trees origin on phenotype.
- **Figure S2:** Predicted phenotypic values as a function of observed ones.
- **Table S2:** Climatic variables description.
- **Figure S3:** Correlation between phenotypes and climatic variables.
- **Table S3:** Population definition.
- **Figure S4:** Allele frequencies as a function of climatic variables.
- **Table S4:** Candidate SNPs associated either to environment variables or phenotypic traits.
- **Table S5:** Maximum overlap between genes involved in response to different climatic variables (**A**) or categories of variables (**B**).
- **Figure S5:** Network (shared names) of enriched biological processes gene ontology terms for transcripts detected as responding to climatic factors (**A**) or phenotypic variables (**B**).
- **Figure S6:** Relationships between trait values and between SNP allelic effect sizes.
- **Supplementary files 1:** SNPs annotations and Gene Ontology term.

