## Supplementary material for "Assessing the potential for assisted gene flow using past introduction of Norway spruce in Southern Sweden: Local adaptation and genetic basis of quantitative traits in trees"

### Supporting Information

**Table S1. Trees origin from available records.**

| Origin | Domain | Country of Origin | Number of trees |
|---|---|---|---|
| Swedish Breeding Programs | Alpine | Germany | 26 |
| | Carpathian | Romania | 84 |
| | Fennoscandia | Sweden | 208 |
| | | Denmark | 35 |
| | Visegrad | Czech Republic | 16 |
| | | Poland | 152 |
| | | Slovakia | 126 |
| | Russia-Baltics | Byelorussia | 255 |
| | | Estonia | 1 |
| | | Lithuania | 10 |
| | | Russia | 2 |
| | Unknown | - | 560 |
| | | Sub-total | 1475 |
| Natural Populations | Alpine | Germany | 9 |
| | | Switzerland | 2 |
| | Carpathian | Romania | 5 |
| | Fennoscandia | Finland | 13 |
| | | Sweden | 16 |
| | Russia-Baltics | Latvia | 4 |
| | | Lithuania | 2 |
| | | Russia | 17 |
| | | Byelorussia | 2 |
| | | Sub-total | 70 |
| | | Total | 1545 |

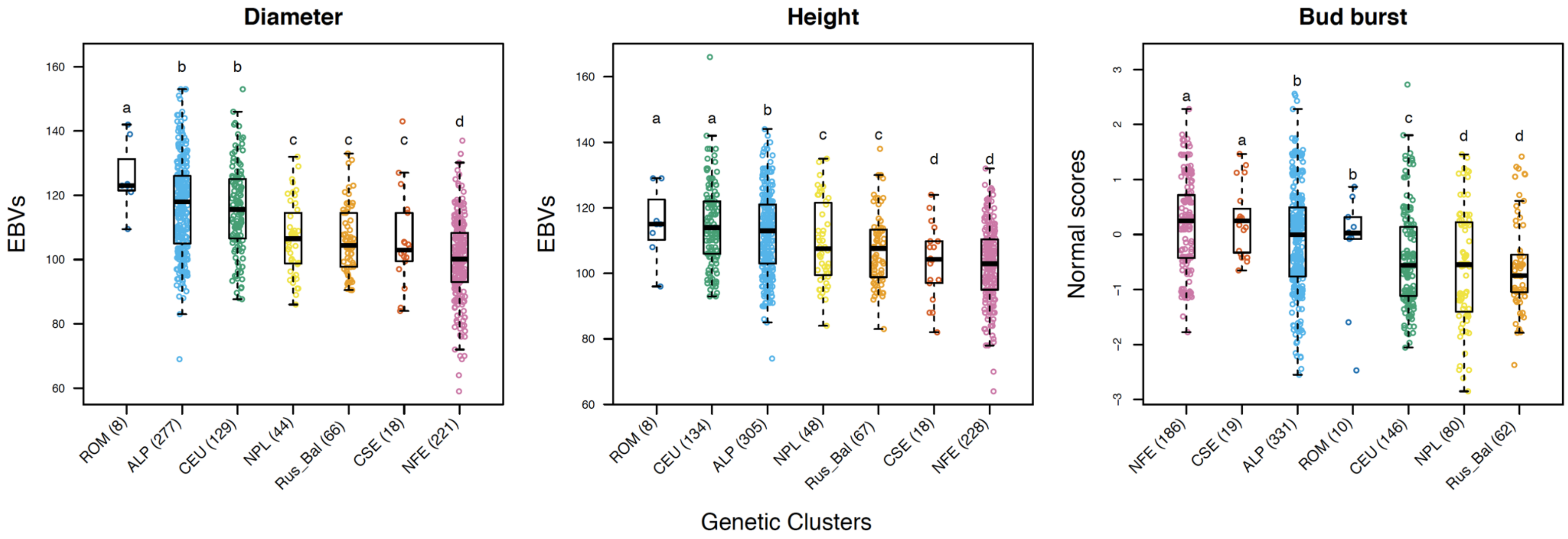


**Figure S1: Influence of trees origin on phenotype.** Diameter, height (breeding values, EBVs) and bud burst (normal-scores) values are represented for the different genetic clusters (Carpathian, dark blue, ROM; Alpine, light blue, ALP; Central Europe, green, CEU; Northern Poland, yellow, NPL; Russia-Baltic, orange, Rus-Bal; Central and Southern Sweden, red, CSE; Fennoscandian, pink, NFE). The number of trees belonging to each genetic cluster is given within parentheses. Letters represent the levels of significance.

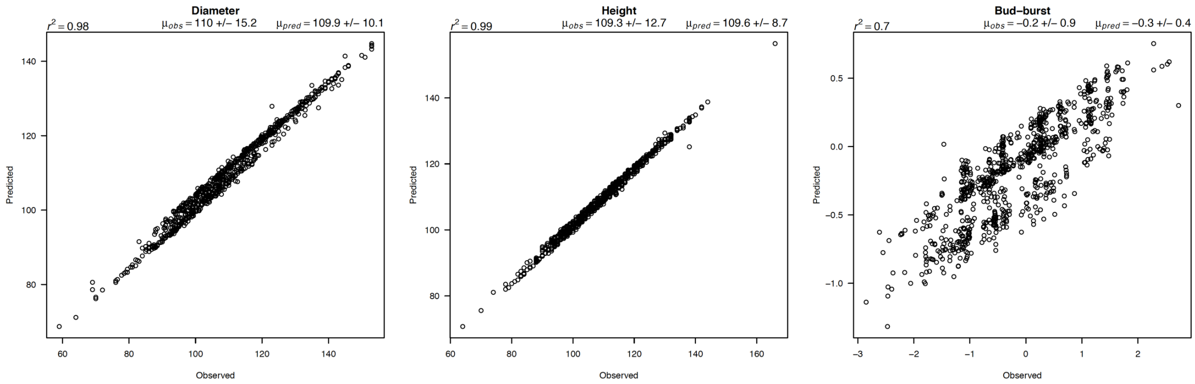


**Figure S2: Predicted phenotypic values as a function of observed ones.**

**Table S2: Climatic variables description**

| Name | Abbrev. | Description |
|---|---|---|
| Temp_1 | µATemp | Annual Mean Temperature |
| Temp_2 | µRangeDuir | Mean Diurnal Range (Mean of monthly (max - min temp)) |
| Temp_3 | MaxTemWarm_Month | Max Temperature of Warmest Month |
| Temp_4 | MinTempCold_Month | Min Temperature of Coldest Month |
| Temp_5 | ARangeTemp | Temperature Annual Range (Temp_3 - Temp_4) |
| Temp_6 | µTempWet_Quart | Mean Temperature of Wettest Quarter |
| Temp_7 | µTempDry_Quart | Mean Temperature of Driest Quarter |
| Temp_8 | µTempWarm_Quart | Mean Temperature of Warmest Quarter |
| Temp_9 | µTempCold_Quart | Mean Temperature of Coldest Quarter |
| Temp_10 | TempSeas | Temperature Seasonality (standard deviation *100) |
| Prec_1 | PrecSeas | Precipitation Seasonality (Coefficient of Variation) |
| Prec_2 | uAPrec | Annual Precipitation |
| Prec_3 | totPrecWet_Month | Precipitation of Wettest Month |
| Prec_4 | totPrecDry_Month | Precipitation of Driest Month |
| Prec_5 | totPrecWet_Quart | Precipitation of Wettest Quarter |
| Prec_6 | totPrecDry_Quart | Precipitation of Driest Quarter |
| Prec_7 | totPrecWarm_Quart | Precipitation of Warmest Quarter |
| Prec_8 | totPrecCold_Quart | Precipitation of Coldest Quarter |
| Moist_1 | AHM | Annual heat-moisture index (Temp_1) / (Prec_2/100) |
| Moist_2 | SHM | Summer heat-moisture index (Temp_3) /(Prec_7/100) |
| Photo_1 | *∆*DL | Average day length in June – in January |

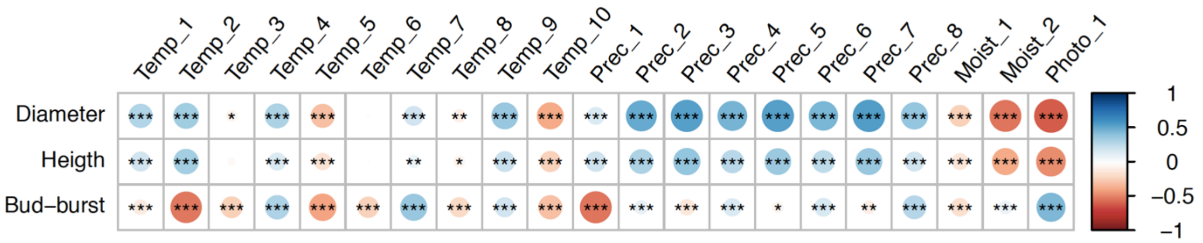


**Figure S3: Correlation between phenotypes and climatic variables.** For each pair of phenotype and climatic variable, the disc diameter represents Pearson's product moment correlation coefficient, *r*, (blue means positive and red negative values). The level of significance is also indicated: ***, $p < 0.001$; **, $p < 0.01$; *, $p < 0.05$. See table S2 above, for variable description.

**Table S3: Population definition**

| Genetic Cluster | Pop# | Size | Latitude | Longitude |
|---|---|---|---|---|
| CSE | 1 | 6 | 55.50 | 13.50 |
| NFE | 2 | 9 | 57.17 | 16.03 |
| | 3 | 6 | 62.40 | 24.29 |
| | 4 | 16 | 61.55 | 12.75 |
| | 5 | 7 | 61.57 | 29.22 |
| | 6 | 11 | 55.50 | 13.50 |
| | 7 | 10 | 58.50 | 13.50 |
| | 8 | 13 | 58.50 | 16.50 |
| | 9 | 8 | 61.50 | 13.50 |
| | 10 | 11 | 61.50 | 16.50 |
| Rus_Bal | 11 | 12 | 54.69 | 25.28 |
| | 12 | 6 | 53.90 | 27.56 |
| | 13 | 23 | 55.18 | 30.17 |
| | 14 | 113 | 53.90 | 27.57 |
| | 15 | 9 | 53.92 | 25.83 |
| | 16 | 96 | 55.58 | 28.18 |
| | 17 | 17 | 53.30 | 34.30 |
| | 18 | 5 | 55.50 | 28.77 |
| | 19 | 7 | 55.50 | 28.50 |
| CEU | 20 | 14 | 49.88 | 19.17 |
| | 21 | 64 | 49.58 | 18.83 |
| | 22 | 9 | 49.56 | 18.89 |
| | 23 | 8 | 50.34 | 22.58 |
| | 24 | 15 | 48.73 | 19.15 |
| | 25 | 27 | 48.80 | 19.64 |
| | 26 | 13 | 50.11 | 12.81 |
| | 27 | 19 | 49.22 | 18.74 |
| | 28 | 13 | 48.75 | 19.67 |
| | 29 | 12 | 49.39 | 19.30 |
| | 30 | 9 | 48.90 | 19.73 |
| | 31 | 5 | 48.80 | 20.67 |
| | 32 | 7 | 49.35 | 19.25 |
| | 33 | 9 | 49.45 | 19.25 |
| | 34 | 7 | 49.50 | 19.50 |
| NPL | 35 | 7 | 52.70 | 23.87 |
| | 36 | 7 | 53.13 | 23.17 |
| | 37 | 5 | 53.41 | 20.34 |
| | 38 | 7 | 52.42 | 23.46 |
| | 39 | 16 | 54.03 | 23.03 |
| ALP | 40 | 13 | 47.63 | 9.64 |
| | 41 | 13 | 48.28 | 8.19 |
| | 42 | 9 | 47.00 | 12.23 |
| ROM | 43 | 9 | 47.24 | 25.70 |
| | 44 | 18 | 47.31 | 24.65 |
| | 45 | 21 | 47.58 | 25.57 |
| | 46 | 10 | 46.36 | 23.05 |
| | 47 | 21 | 46.92 | 25.35 |
| | 48 | 5 | 46.82 | 25.12 |
| **Population definition for trees with inferred original location (Fig. S4)** | | | | |
| CSE | 49 | 21 | 57.01 | 14.36 |
| NFE | 50 | 195 | 59.64 | 16.33 |
| Rus_Bal | 51 | 38 | 54.62 | 28.30 |
| CEU | 52 | 53 | 49.38 | 19.01 |
| NPL | 53 | 28 | 52.98 | 22.42 |
| ALP | 54 | 267 | 47.63 | 9.64 |
| | 55 | 27 | 47.63 | 9.64 |
| | 56 | 48 | 47.63 | 9.64 |
| ROM | 57 | 17 | 47.07 | 25.00 |

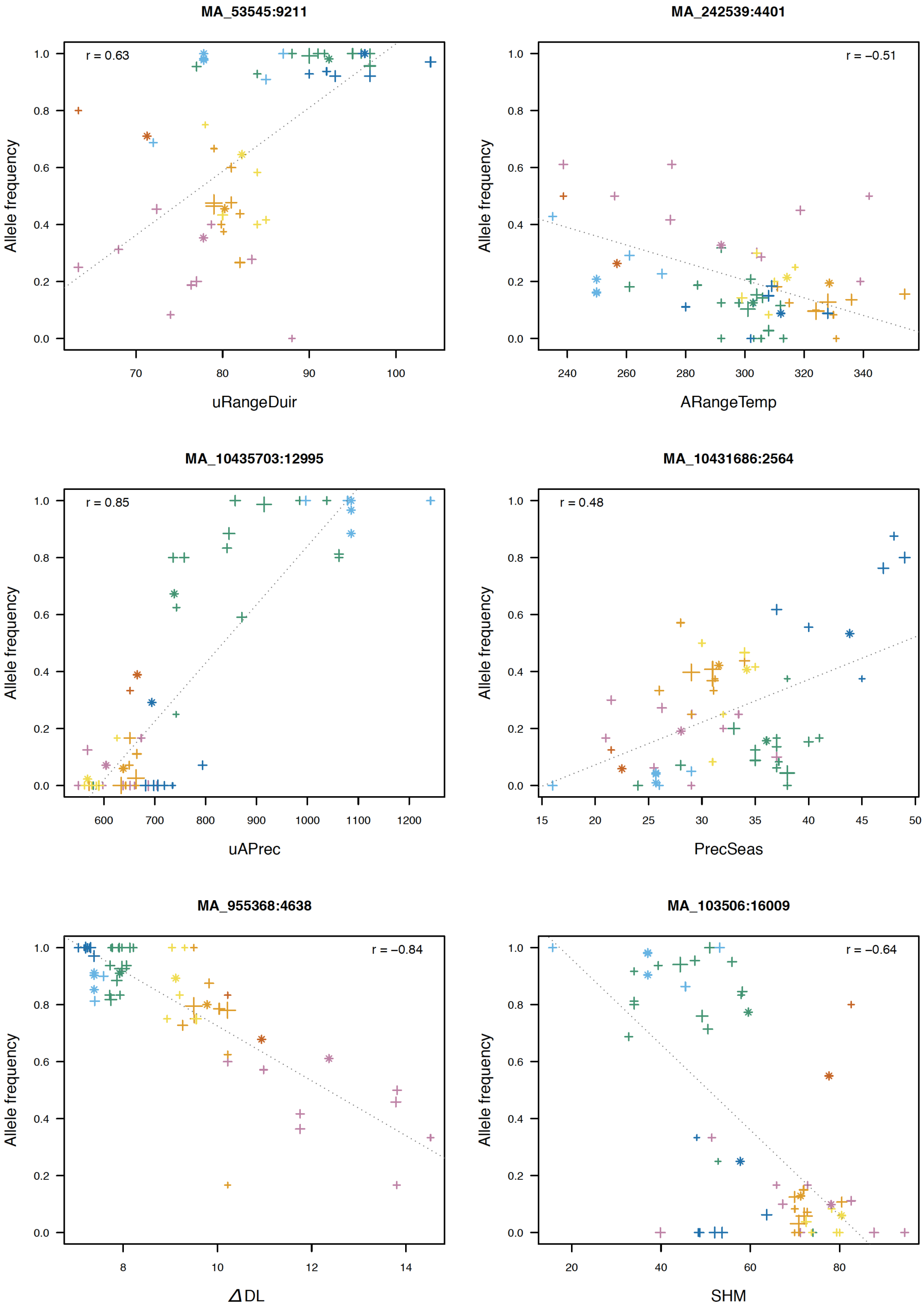


**Figure S4: Allele frequencies as a function of climatic variables.** The allele frequencies of the SNPs having the highest Bayes Factor (Bayenv2) among the candidate SNPs are represented as a function of the corresponding climatic variable. For each graph, the main title corresponds to the SNP name (scaffold:position), the dotted line is the regression between both factors, and *r*, is the Pearson's correlation coefficient. "Plus" signs are allele frequencies in each populations: the bigger it is, the larger the number of individuals in the population. Stars represent allele frequencies for individuals with inferred original location. Finally, colors represent the different genetic clusters (Carpathian, dark blue, ROM; Alpine, light blue, ALP; Central Europe, green, CEU; Northern Poland, yellow, NPL; Russia-Baltic, orange, Rus-Bal; Central and Southern Sweden, red, CSE; Fennoscandian, pink, NFE).

**Table S4: Candidate SNPs associated either to environment variables or phenotypic traits.**

| Climatic var. | BF | N | Transc. | Redundancy | Intergenic | Intron | Exon | NSyn | Syn |
|---|---|---|---|---|---|---|---|---|---|
| Temp_1 | 241 | 46 | 43 | 7 | 22 | 54 | 24 | 83 | 17 |
| Temp_10 | 257 | 91 | 66 | 27 | 12 | 54 | 34 | 62 | 38 |
| Temp_2 | 257 | 63 | 60 | 5 | 22 | 38 | 40 | 63 | 35 |
| Temp_3 | 257 | 58 | 51 | 12 | 21 | 41 | 38 | 55 | 45 |
| Temp_4 | 257 | 75 | 58 | 23 | 21 | 44 | 35 | 80 | 20 |
| Temp_5 | 257 | 85 | 75 | 12 | 15 | 44 | 41 | 63 | 37 |
| Temp_6 | 257 | 76 | 61 | 20 | 18 | 45 | 37 | 70 | 30 |
| Temp_7 | 257 | 68 | 59 | 13 | 12 | 47 | 41 | 61 | 39 |
| Temp_8 | 257 | 50 | 50 | 0 | 24 | 42 | 34 | 41 | 59 |
| Temp_9 | 257 | 162 | 99 | 39 | 15 | 49 | 36 | 64 | 39 |
| Prec_1 | 257 | 65 | 59 | 9 | 18 | 43 | 38 | 82 | 21 |
| Prec_2 | 362 | 277 | 98 | 65 | 19 | 53 | 29 | 59 | 38 |
| Prec_3 | 257 | 143 | 69 | 52 | 20 | 45 | 34 | 59 | 41 |
| Prec_4 | 434 | 344 | 128 | 63 | 18 | 52 | 30 | 60 | 40 |
| Prec_5 | 257 | 139 | 68 | 51 | 19 | 45 | 36 | 61 | 39 |
| Prec_6 | 396 | 305 | 112 | 63 | 19 | 52 | 29 | 59 | 41 |
| Prec_7 | 257 | 148 | 70 | 53 | 20 | 45 | 35 | 60 | 40 |
| Prec_8 | 352 | 268 | 95 | 65 | 18 | 51 | 31 | 58 | 42 |
| Moist_1 | 257 | 156 | 72 | 54 | 19 | 44 | 37 | 59 | 41 |
| Moist_2 | 257 | 126 | 63 | 50 | 21 | 42 | 37 | 57 | 46 |
| Photo_1 | 864 | 230 | 148 | 36 | 17 | 48 | 35 | 63 | 37 |
| **Phenotype** | | ***s* < 0.1** | **Transc.** | **Redundancy** | **Intergenic** | **Intron** | **Exon** | **NSyn** | **Syn** |
| Bud-burst | | 32 | 15 | 53 | 6 | 56 | 38 | 58 | 41 |
| Diameter | | 180 | 131 | 27 | 15 | 51 | 34 | 62 | 38 |
| Height | | 175 | 138 | 21 | 17 | 46 | 37 | 62 | 39 |

For each climatic and phenotypic variable, the number of significant SNPs is reported. *BF* gives the number of SNPs having either a Bayes Factor > 150 or a Bayes Factor > 20 and within the 0.1% highest BF. *N* is the number of those SNPs within the top 1% of Spearman's correlation coefficients. *s* is the analogue of *q*-value for *false sign rate* detection (Stephens 2017). Details of climatic variables are given in table S2. Transcript (Transc.) is the number of unique transcript (defined as the closest transcript to each SNPs position) and redundancy is the number of candidate SNPs over the number of unique transcript. The columns Intergenic, Intron, Exon, NSyn (non-synonymous sites) and Syn (synonymous sites) give the percentage of SNPs belonging to each of these categories.

**Table S5A: Maximum overlap (%) between genes involved in response to different climatic variables.**

| | Temp_1 | Temp_2 | Temp_3 | Temp_4 | Temp_5 | Temp_6 | Temp_7 | Temp_8 | Temp_9 | Temp_10 | Prec_1 | Prec_2 | Prec_3 | Prec_4 | Prec_5 | Prec_6 | Prec_7 | Prec_8 | Moist_1 | Moist_2 |
|---|---|---|---|---|---|---|---|---|---|---|---|---|---|---|---|---|---|---|---|---|
| **Temp_2** | 0 | - | - | - | - | - | - | - | - | - | - | - | - | - | - | - | - | - | - | - |
| **Temp_3** | 23 | 0 | - | - | - | - | - | - | - | - | - | - | - | - | - | - | - | - | - | - |
| **Temp_4** | 5 | 4 | 4 | - | - | - | - | - | - | - | - | - | - | - | - | - | - | - | - | - |
| **Temp_5** | 0 | 2 | 5 | 6 | - | - | - | - | - | - | - | - | - | - | - | - | - | - | - | - |
| **Temp_6** | 14 | 0 | 37 | 4 | 7 | - | - | - | - | - | - | - | - | - | - | - | - | - | - | - |
| **Temp_7** | 2 | 5 | 3 | 31 | 5 | 10 | - | - | - | - | - | - | - | - | - | - | - | - | - | - |
| **Temp_8** | 12 | 2 | 4 | 52 | 0 | 10 | 32 | - | - | - | - | - | - | - | - | - | - | - | - | - |
| **Temp_9** | 33 | 0 | 54 | 4 | 3 | 47 | 3 | 4 | - | - | - | - | - | - | - | - | - | - | - | - |
| **Temp_10** | 0 | 0 | 7 | 6 | 43 | 13 | 20 | 6 | 10 | - | - | - | - | - | - | - | - | - | - | - |
| **Prec_1** | 0 | 5 | 0 | 2 | 2 | 0 | 2 | 0 | 2 | 3 | - | - | - | - | - | - | - | - | - | - |
| **Prec_2** | 2 | 0 | 2 | 2 | 3 | 1 | 3 | 0 | 8 | 30 | 7 | - | - | - | - | - | - | - | - | - |
| **Prec_3** | 2 | 8 | 0 | 0 | 2 | 0 | 2 | 0 | 8 | 22 | 2 | 43 | - | - | - | - | - | - | - | - |
| **Prec_4** | 2 | 0 | 3 | 4 | 7 | 4 | 5 | 2 | 10 | 35 | 10 | 69 | 42 | - | - | - | - | - | - | - |
| **Prec_5** | 2 | 9 | 0 | 0 | 2 | 0 | 0 | 0 | 8 | 24 | 2 | 43 | 75 | 40 | - | - | - | - | - | - |
| **Prec_6** | 2 | 0 | 2 | 4 | 3 | 1 | 3 | 2 | 8 | 31 | 8 | 69 | 42 | 89 | 40 | - | - | - | - | - |
| **Prec_7** | 2 | 6 | 0 | 0 | 2 | 0 | 2 | 0 | 8 | 27 | 2 | 51 | 72 | 47 | 82 | 47 | - | - | - | - |
| **Prec_8** | 0 | 0 | 3 | 2 | 9 | 4 | 5 | 0 | 8 | 35 | 10 | 58 | 39 | 81 | 38 | 78 | 46 | - | - | - |
| **Moist_1** | 9 | 0 | 7 | 4 | 2 | 6 | 3 | 2 | 17 | 25 | 2 | 53 | 43 | 50 | 41 | 49 | 49 | 44 | - | - |
| **Moist_2** | 12 | 6 | 10 | 2 | 2 | 11 | 2 | 2 | 25 | 19 | 2 | 32 | 48 | 32 | 51 | 32 | 52 | 30 | 51 | - |
| **Photo_1** | 0 | 18 | 0 | 2 | 2 | 1 | 2 | 0 | 0 | 1 | 10 | 2 | 6 | 2 | 6 | 2 | 4 | 4 | 0 | 2 |

**Table S5B: Average overlap (%).**

| | Temperature | Precipitation | Moisture |
|---|---|---|---|
| **Temperature** | 12 | - | - |
| **Precipitation** | 5 | 44 | - |
| **Moisture** | 8 | 38 | 51 |
| **Photoperiod** | 3 | 5 | 1 |

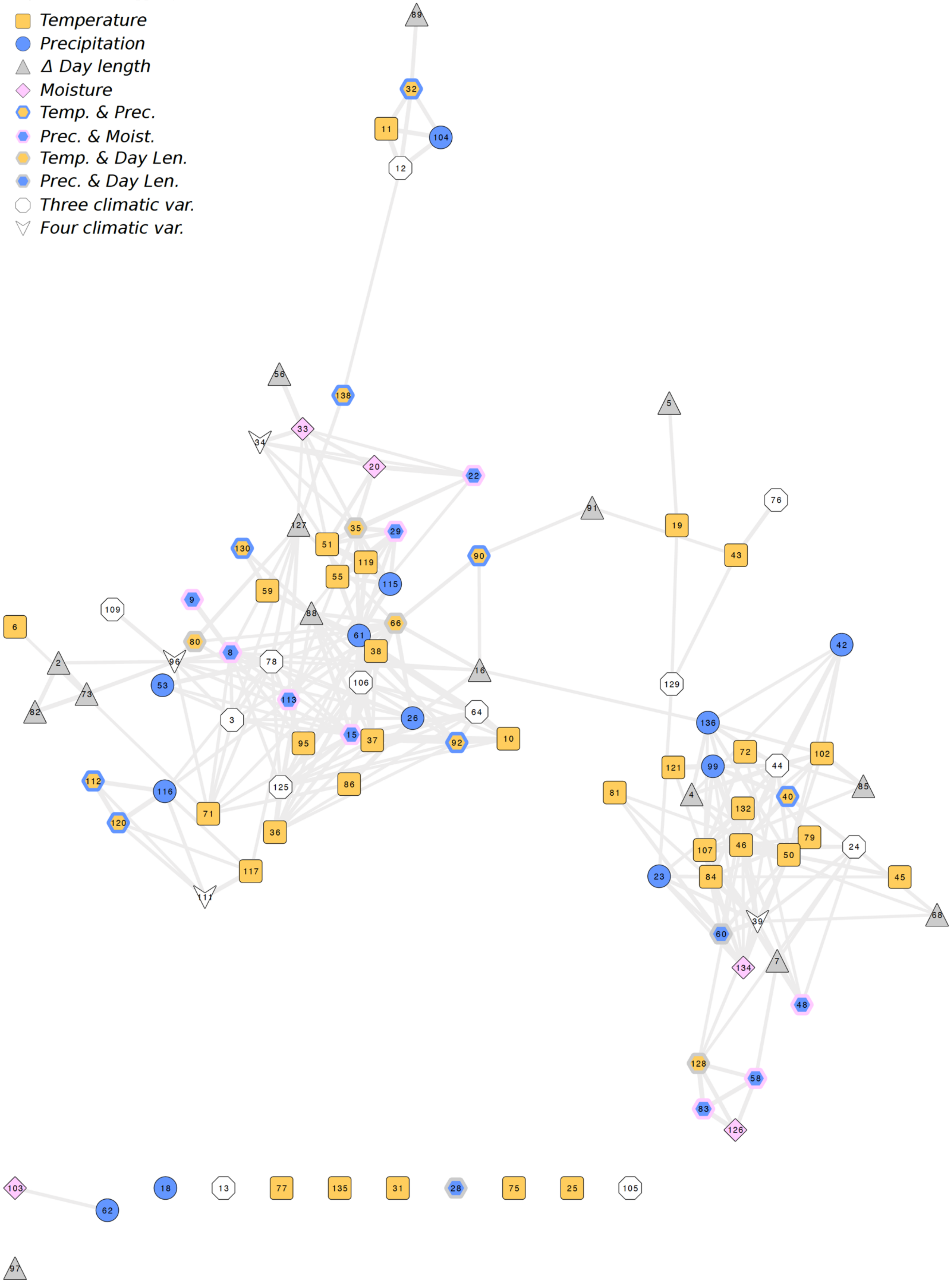


1 DNA duplex unwinding
2 Golgi localization
3 Golgi organization
4 Lewis a epitope biosynthesis
5 RNA metabolism
6 RNA-dependent DNA biosynthesis
7 acetyl-CoA biosynthesis
8 actin filament bundle assembly
9 actin filament severing
10 adventitious root development
11 auxin metabolism
12 auxin polar transport

13 calcium ion transport
14 carpel development
15 cell morphogenesis
16 cell plate formation involved in plant-type cell wall biogenesis
17 cell wall pectin biosynthesis
18 cellular component organization or biogenesis
19 cellular macromolecule biosynthesis
20 cellular response to nutrient levels
21 cellular response to salt stress
22 cellular response to sulfur starvation
23 ceramide metabolism
24 chlorophyll catabolism
25 chloroplast RNA modification
26 chloroplast fission
27 chromatin remodeling
28 chromosome segregation
29 cold acclimation
30 covalent chromatin modification
31 cullin deneddylation
32 cytokinin catabolism
33 defense response to fungus, incompatible interaction
34 defense response to insect
35 detection of temperature stimulus
36 embryo sac cellularization
37 embryonic pattern specification
38 endosperm development
39 fatty acid beta-oxidation
40 flavonol biosynthesis
41 floral organ formation
42 fucose biosynthesis
43 gene silencing by miRNA
44 gluconeogenesis
45 glyceraldehyde-3-phosphate biosynthesis
46 glycerol catabolism
47 glycolytic process
48 glycoside catabolism
49 glycosylceramide catabolism
50 glyoxylate cycle
51 gravitropism
52 heterochromatin assembly
53 histone H3-K36 methylation
54 homogalacturonan biosynthesis
55 hyperosmotic response
56 immune response
57 indeterminate inflorescence morphogenesis
58 indole glucosinolate biosynthesis
59 interspecies interaction between organisms
60 long-chain fatty acid metabolism
61 long-day photoperiodism, flowering
62 mRNA polyadenylation
63 maintenance of meristem identity
64 maintenance of root meristem identity
65 maintenance of shoot apical meristem identity
66 meiotic mismatch repair
67 meristem maintenance
68 miRNA catabolism
69 microsporocyte differentiation
70 microtubule cytoskeleton organization
71 mitochondrial DNA replication
72 mitochondrial electron transport, cytochrome c to oxygen
73 mitochondrion localization
74 mitotic G2 phase
75 mitotic interphase
76 negative regulation of programmed cell death
77 nitrogen compound metabolism
78 nucleosome assembly
79 nucleotide phosphorylation
80 nucleotide-excision repair, DNA incision, 5'-to lesion
81 para-aminobenzoic acid metabolism
82 peroxisome localization
83 phenylpropanoid metabolism
84 phosphatidylinositol biosynthesis
85 photosystem II repair
86 plastid DNA replication
87 pollen germination
88 pollen tube development
89 polyamine biosynthesis
90 positive regulation of cell cycle
91 positive regulation of protein geranylgeranylation
92 primary root development
93 primary shoot apical meristem specification
94 production of small RNA involved in gene silencing by RNA
95 proteasome core complex assembly
96 protein import into peroxisome matrix
97 protein processing
98 protein targeting to membrane
99 pyridoxine biosynthesis
100 red light signaling pathway
101 red, far-red light phototransduction
102 reductive pentose-phosphate cycle
103 regulation of RNA splicing
104 regulation of auxin biosynthesis
105 regulation of circadian rhythm
106 regulation of flower development
107 regulation of glycolytic process
108 regulation of ion transport
109 regulation of protein localization
110 resolution of meiotic recombination intermediates
111 response to auxin
112 response to cobalt ion
113 response to endoplasmic reticulum stress
114 response to far red light
115 response to high light intensity
116 response to hydrogen peroxide
117 response to misfolded protein
118 response to radiation
119 response to temperature stimulus
120 response to zinc ion
121 riboflavin biosynthesis
122 root hair elongation
123 root meristem specification
124 specification of floral organ identity
125 sporopollenin biosynthesis
126 suberin biosynthesis
127 telomere maintenance in response to DNA damage
128 toxin catabolism
129 translational initiation
130 transport of virus in host, cell to cell
131 triglyceride biosynthesis
132 triglyceride mobilization
133 tryptophan biosynthesis
134 very long-chain fatty acid biosynthesis
135 water transport
136 xyloglucan biosynthesis
137 zinc II ion transport
138 zinc ion homeostasis

**Figure S5A: Network (shared names) of enriched biological processes gene ontology terms for transcripts detected as responding to climatic factors.**

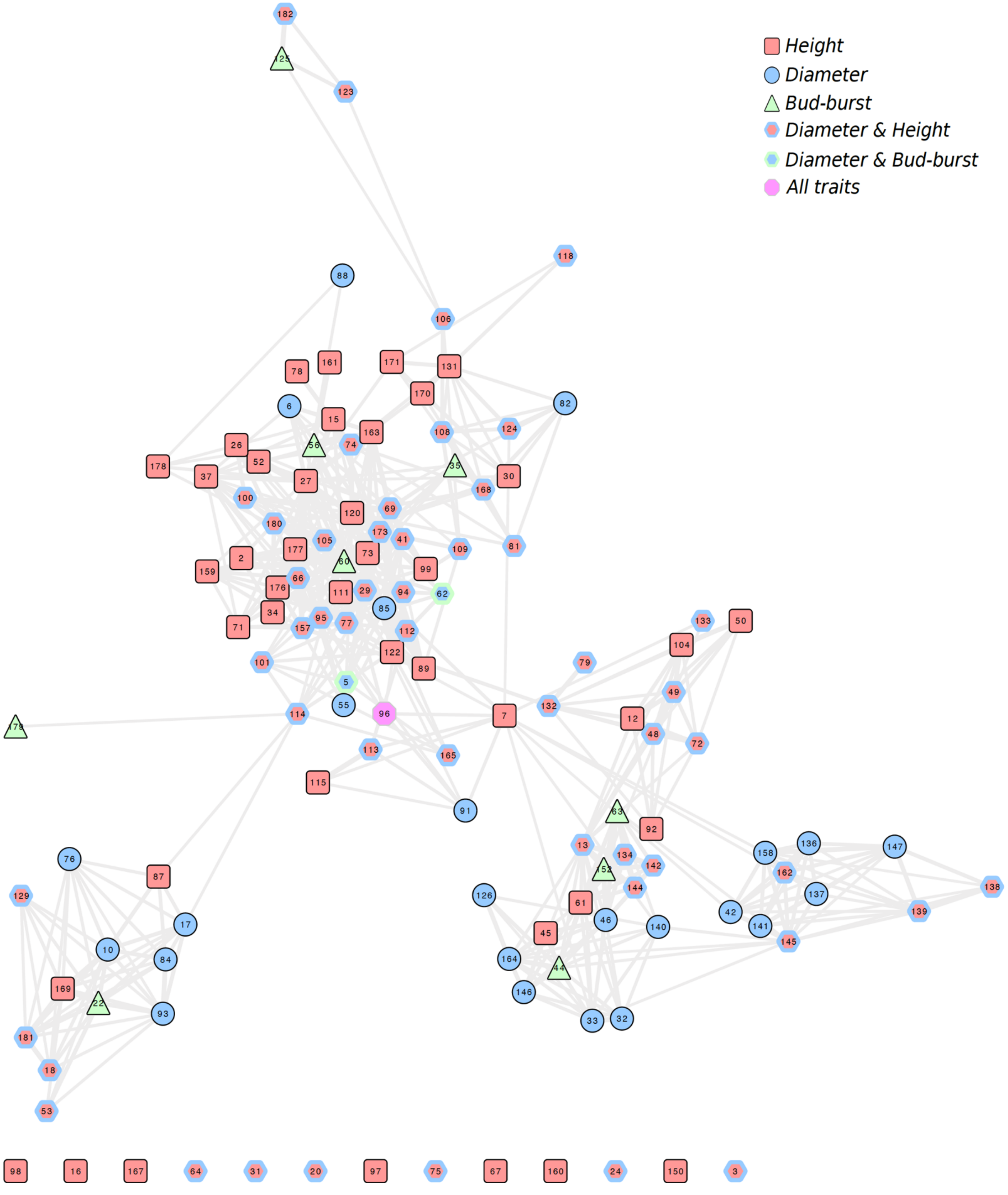


1 ATP biosynthesis
2 D-ribose metabolism
3 DNA endoreduplication
4 ER to Golgi vesicle-mediated transport
5 Golgi organization
6 L-serine metabolism
7 MAPK cascade
8 amino acid transport
9 ammonium transmembrane transport
10 ammonium transport
11 anatomical structure morphogenesis
12 organ morphogenesis
13 anthocyanin accumulation in tissues in response to UV light
14 aromatic amino acid family biosynthesis
15 aromatic amino acid family metabolism
16 asymmetric cell division
17 basic amino acid transport
18 cadmium ion transport
19 calcium ion transport
20 carbohydrate metabolism
21 carboxylic acid metabolism
22 cation transport
23 cell differentiation
24 cell proliferation
25 cell tip growth
26 cell wall modification
27 cellular amino acid biosynthesis
28 cellular cation homeostasis
29 cellular lipid catabolism
30 cellular macromolecule biosynthesis
31 cellular process
32 cellular response to iron ion starvation

33 cellular response to phosphate starvation
34 cellulose biosynthesis
35 chlorophyll biosynthesis
36 chloroplast relocation
37 chorismate biosynthesis
38 coenzyme biosynthesis
39 coumarin biosynthesis
40 cysteine biosynthesis
41 cytochrome complex assembly
42 cytokinin-activated signaling pathway
43 defense response to bacterium
44 defense response to fungus
45 detection of biotic stimulus
46 detection of temperature stimulus
47 divalent metal ion transport
48 embryo development
49 embryo development ending in seed dormancy
50 embryo sac egg cell differentiation
51 endonucleolytic cleavage in ITS1
52 endonucleolytic cleavage
53 establishment of localization
54 fatty acid beta-oxidation
55 galactolipid biosynthesis
56 glucosinolate biosynthesis
57 glucuronoxylan metabolism
58 glycine catabolism
59 glycine metabolism
60 glycolytic process
61 gravitropism
62 hydrogen peroxide catabolism
63 hyperosmotic response
64 indole-containing compound metabolism
65 indoleacetic acid biosynthesis
66 inositol biosynthesis
67 intra-Golgi vesicle-mediated transport
68 iron ion transport
69 iron-sulfur cluster assembly
70 isopentenyl diphosphate biosynthesis, methylerythritol 4-phosphate pathway
71 jasmonic acid biosynthesis
72 leaf morphogenesis
73 lipoate metabolism
74 mRNA splicing, via spliceosome
75 metabolism
76 methylammonium transport
77 methylglyoxal catabolism to D-lactate via S-lactoyl-glutathione
78 mitochondrial mRNA modification
79 multidimensional cell growth
80 myo-inositol hexakisphosphate biosynthesis
81 ncRNA metabolism
82 negative regulation of transcription, DNA-templated
83 negative regulation of translation
84 nitrate transport
85 nucleosome assembly
86 nucleotide transport
87 nucleotide-sugar transport
88 one-carbon metabolism
89 para-aminobenzoic acid metabolism
90 pentose-phosphate shunt
91 peptidyl-tyrosine dephosphorylation
92 phloem or xylem histogenesis
93 phosphate ion transport
94 phosphatidylglycerol biosynthesis
95 phospholipid biosynthesis
96 phosphorylation
97 photorespiration
98 photosynthesis
99 photosystem II assembly
100 plant-type secondary cell wall biogenesis
101 plastid organization
102 pollen development
103 pollen exine formation
104 pollen sperm cell differentiation
105 polysaccharide biosynthesis
106 positive regulation of auxin metabolism
107 positive regulation of organelle organization
108 positive regulation of transcription, DNA-templated
109 positive regulation of tryptophan metabolism
110 proteasome assembly
111 proteasome core complex assembly
112 proteasome-mediated ubiquitin-dependent protein catabolism
113 protein deubiquitination
114 protein import into peroxisome matrix
115 protein sumoylation
116 protein targeting to membrane
117 protein ubiquitination
118 proteolysis
119 purine nucleotide biosynthesis
120 rRNA processing
121 regulation of cellular metabolism
122 regulation of chromosome organization
123 regulation of hormone levels
124 regulation of macromolecule metabolism
125 regulation of pH
126 regulation of plant-type hypersensitive response
127 regulation of primary metabolism
128 regulation of protein dephosphorylation
129 regulation of proton transport
130 regulation of response to biotic stimulus
131 regulation of translation
132 regulation of unidimensional cell growth
133 response to abiotic stimulus
134 response to blue light
135 response to cadmium ion
136 response to carbon dioxide
137 response to chitin
138 response to cobalt ion
139 response to copper ion
140 response to endoplasmic reticulum stress
141 response to ethylene
142 response to far red light
143 response to heat
144 response to high light intensity
145 response to misfolded protein
146 response to nematode
147 response to nitrate
148 response to red light
149 response to salt stress
150 response to stimulus
151 response to sucrose
152 response to temperature stimulus
153 response to zinc ion
154 root development
155 root hair elongation
156 root morphogenesis
157 salicylic acid biosynthesis
158 salicylic acid mediated signaling pathway
159 shikimate biosynthesis
160 single organismal cell-cell adhesion
161 suberin biosynthesis
162 sugar mediated signaling pathway
163 sulfur amino acid metabolism
164 systemic acquired resistance
165 thylakoid membrane organization
166 tissue development
167 transcription from plastid promoter
168 transcription, DNA-templated
169 transition metal ion transport
170 translational elongation
171 translational initiation
172 transport
173 tryptophan catabolism
174 ubiquitin-dependent protein catabolism
175 unidimensional cell growth
176 unsaturated fatty acid biosynthesis
177 very long-chain fatty acid biosynthesis
178 vitamin metabolism
179 water transport
180 xylan biosynthesis
181 zinc II ion transport
182 zinc ion homeostasis

**Figure S5B: Network (shared names) of enriched biological processes gene ontology terms for transcripts detected as involved in the control of phenotypic traits.**

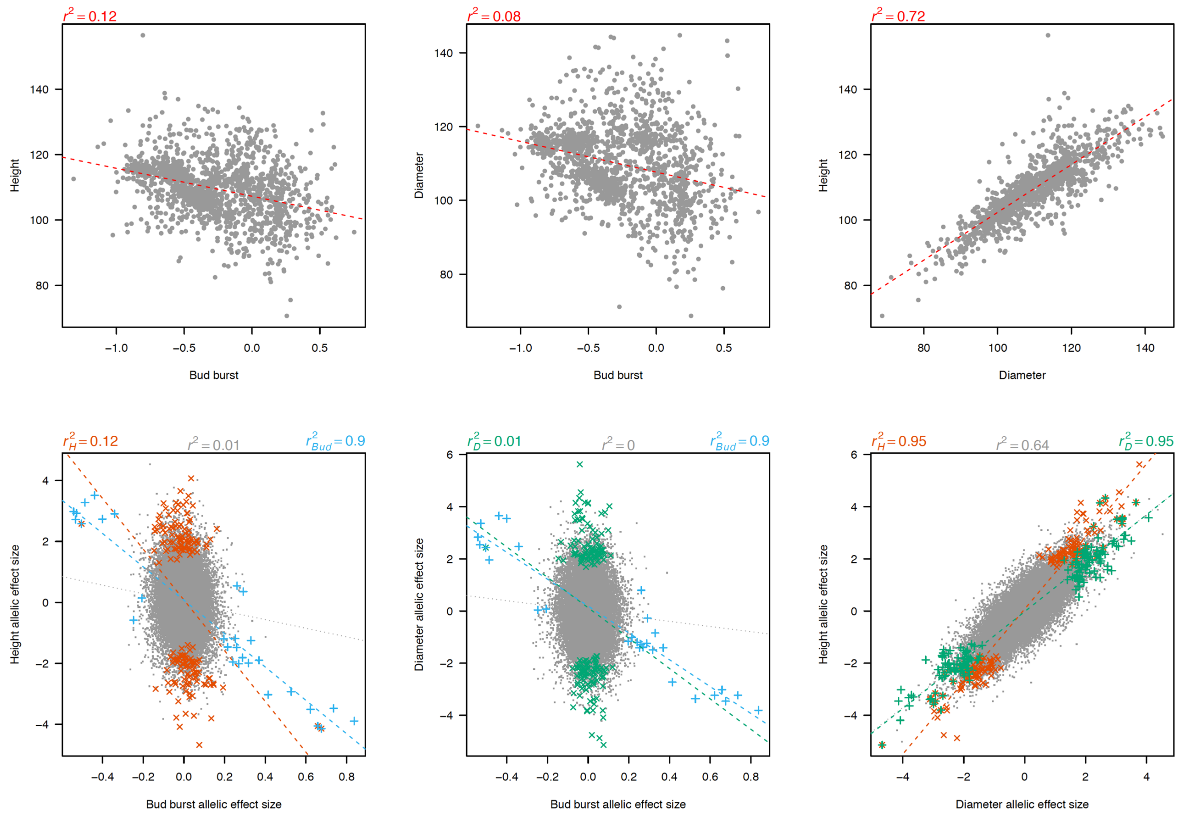


**Figure S6: Relationships between trait values (upper panels) and between SNP allelic effect sizes (lower panels).**

Top panels represent *height* as a function of *bud-burst* (left), *diameter* as a function of *bud-burst* (middle) and *height* as a function of *diameter* (right), the dotted lines are the linear regression between trait values, adjusted R-squared are indicated ($r^2$). Bottom panels represent the same relationships than top panels but for SNP allelic effect sizes. For each pair of traits, three linear regressions are represented (dotted lines) between all SNPs (grey), or between SNPs detected as significantly affecting height (orange), bud-burst (blue) or diameter (green), corresponding $r^2$ are provided.
